# Stimulation of the final cell cycle in the stomatal lineage by the cyclin CYCD7;1 under regulation of the MYB transcription factor FOUR-LIPS

**DOI:** 10.1101/207845

**Authors:** Farah Patell, David Newman, Eunkyoung Lee, Zidian Xie, Carl Collins, Erich Grotewold, James A.H. Murray, Walter Dewitte

## Abstract

Stomatal guard cells are formed through a sequence of asymmetric and symmetric divisions in the epidermis of the sporophyte of most land plants. We show that several D-type cyclins are consecutively activated in the stomatal linage in the epidermis of *Arabidopsis thaliana*. Whereas *CYCD2;1* and *CYCD3;2* are activated in the meristemoids early in the lineage, *CYCD7;1* is activated before the final division. *CYCD7;1* expression peaks in the guard mother cell, where its transcription is modulated by the FOUR-LIPS/MYB88 transcription factor. FOUR-LIPS/MYB88 interacts with the *CYCD7;1* promoter and represses CYCD7;1 transcription. *CYCD7;1* stimulates the final symmetric division in the stomatal lineage, since guard cell formation is delayed in the *cycd7;1* mutant epidermis and guard mother cell (GMC) divisions in *four-lips* mutant guard mother cells are limited by loss of function of *CYCD7;1*. Hence, the precise activation of a specific D-type cyclin, CYCD7;1, is required for correct timing of the last symmetric division that creates the stomatal guards cells, and *CYCD7;1* expression is regulated by the FLP/MYB pathway that ensures cell cycle arrest in the stomatal guard cells.

**Summary Statement:** The formation of paired guard cells in the epidermis of the *Arabidopsis thaliana* shoot, requires the activity of the D-type cyclin CYCD7;1 for the normal timing of the final division.

## INTRODUCTION

Stomata are micro-valves in the epidermis of the shoot that mediate gas exchange between internal tissues and the atmosphere and consist of two specialized sister guard cells (GCs) surrounding a pore. The evolution of stomata is associated with the colonization of land by early plants (Vaten and Bergmann, 2012). In *Arabidopsis*, the stomatal lineage arises from stem cell-like meristemoid mother cells (Pillitteri and Torii, 2012), which divide asymmetrically to form a meristemoid (M) that develops into an ovoid guard mother cell (GMC). Finally GMCs divide symmetrically, and the daughters differentiate into the two GCs bordering the stomatal pore.

CDK activity regulates progression through the mitotic cell cycle. In addition to the canonical conserved CDK homologous to yeast cdc2+*/*CDC28 and mammalian CDK1, which is called CDKA in plants, plants encode an additional unique type of CDK known as CDKB. Plant cyclins are represented by three major groups, A-, B- and D-types (De Veylder et al., 2007). In the stomatal lineage, the symmetric GMC division requires the cyclin-dependent kinase *CDKB1;1* and cyclin *CYCA2* (Boudolf et al., 2004; Vanneste et al., 2011), after which GCs stop further cell cycle progression, reportedly with a DNA content of 2C indicative of G1 arrest (Melaragno et al., 1993). *FOUR-LIPS* (*FLP*) and the closely related gene *MYB88* are required to prevent multiple symmetric divisions of the GMC through direct regulation of multiple cell cycle-related genes (Xie et al., 2010).

Cyclin D (CYCD)-CDK complexes have a primary role in regulating the G1/S transition through control of RETINOBLASTOMA-RELATED (RBR) protein activity and thereby the activity of various E2F factor complexes (Magyar et al., 2012; Menges et al., 2006; Zhao et al., 2012). In higher plants, the different *CYCDs* can be classified into six conserved subgroups (Menges et al., 2007), and there is accumulating evidence for distinct developmental roles for different CYCD subgroups (Dewitte et al., 2007; Nieuwland et al., 2009; Sanz et al., 2011; Sozzani et al., 2010). Whilst *RBR* has been shown to have a role in regulating asymmetric divisions of the stomatal lineage (Weimer et al., 2012), the role of *CYCD*s as regulators of RBR function in this process has not been investigated. Here we show that *CYCD7;1* is under control of the FLP/MYB88 transcription factors and stimulates the terminal division of the GMC.

## RESULTS AND DISCUSSION

### *CYCD* expression in the stomatal lineage

Given the rate limiting role of several CYCD genes in proliferation of plant tissues (Dewitte et al., 2007), we examined the expression of *CYCD* genes in the stomatal lineage of the *Arabidopsis* shoot with the exception of *CYCD4;2* and *CYCD5;1*. Only *CYCD2;1, CYCD3;2* and *CYCD7;1* reporters showed activity (Fig. 1). *CYCD3;2* expression was activated in meristemoids and gradually declined in GMCs and GCs, becoming undetectable in mature GCs (mGCs; Fig. 1B). *CYCD2;1_pro_∷CYCD2;1-GFP* fluorescence was detected throughout the later lineage in nuclei of meristemoids, GMCs, and young GCs (Fig. 1A). In line with this, the available ATH1 GeneChip data of the stomatal lineage suggest the presence of *CYCD2;1* transcripts throughout the different phases of the stomatal development and a maximum of *CYCD3;2* expression in meristemoids (eFP browser, data source Guard Cell). In contrast, *CYCD7;1_pro_∷GFP* was expressed in late meristemoids (LM), reached its maximum in GMCs, persisted until after the symmetric division and declined as GCs matured. The GFP signal of a fusion protein transgene consisting of the *CYCD7;1* promoter and genomic coding region fused to *eGFP* was relatively weak, and gave strongest detectable signal in LM and early GMC nuclei (Fig. 1C-D), perhaps due to destabilisation of eGFP as a result of fusion to CYCD7;1. This expression pattern was consistent in stomatal lineages in the cotyledon, leaf and stem epidermis (Fig. 1E-F). Interestingly, *CYCD7;1* expression as determined by both the transcriptional reporter and translational *CYCD7;1- GFP* fusion gene was not detected in sporophytic tissue outside the stomatal lineage, although it was observed in the sperm cells of pollen. Given that *CYCD7;1* is not represented on the ATH1 chip, our current knowledge on *CYCD7;1* expression is still fragmented, but these specific expression dynamics of *CYCD2;1*, *CYCD3;2* and *CYCD7;1* are in agreement with the transcript levels in the stomatal lineage detected by Adrian et al, 2015. Nevertheless, we cannot exclude expression of *CYCD7;1* elsewhere under different experimental conditions. These observations suggested roles for *CYCD2;1*, *CYCD3;2* and particularly *CYCD7;1* in the final phases of stomatal development.

**Figure 1.**
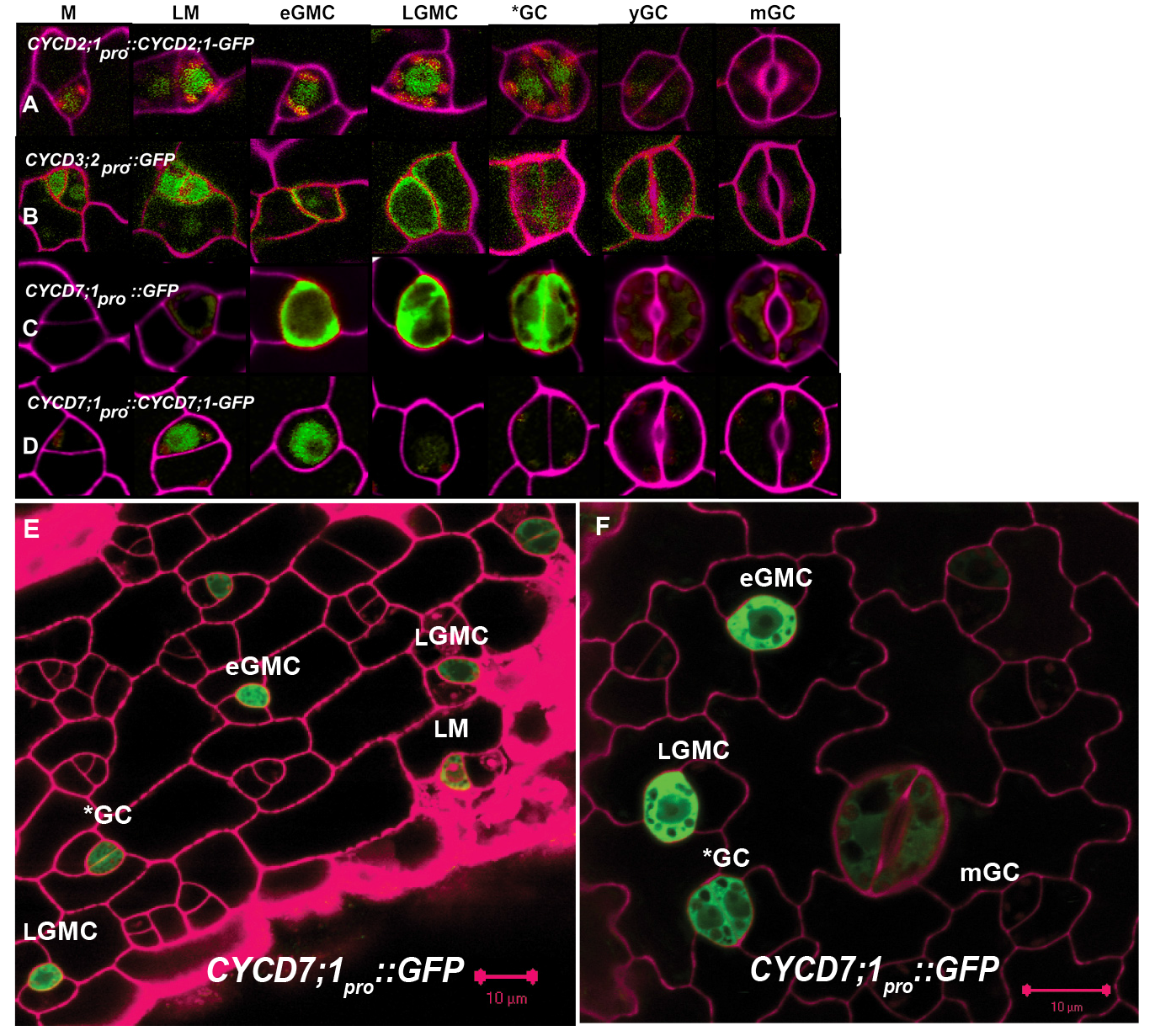
*CYCD* expression in the stomatal lineage. **(A)** Expression of *CYCD2;1;pro∷CYCD2;1-GFP*, **(B)** *CYCD3;2pro∷GFP,* **(C)** *CYCD7;1pro∷GFP*, **(D)** *CYCD7;1pro∷CYCD7;1-GFP* in M (meristemoid), LM (late meristemoid), eGMC (early GMC), LGMC (late GMC), *GC (recently formed GC pair), yGC (young GC pair) and mGC (mature GC). **(E)** and **(F)**. *CYCD7;1pro∷GFP* expression in leaf epidermis showing expression in eGMCs, LGMCs and *GCs

### *CYCD7;1* is transcriptionally controlled by FLP/ MYB88

*FOUR LIPS (FLP)* and its close relative *MYB88* limit GMC division by directly regulating multiple cell cycle genes (Xie et al., 2010), and in *flp* mutants GMC daughter cells reiterate the symmetric division program generating linear clusters of GCs. *FLP* is activated in late GMCs (Xie et al., 2010) coinciding with the timing of reduced CYCD7;1-GFP fluorescence (Fig. 1D). These observations suggested that FLP/MYB88 could negatively regulate *CYCD7;1*, so we examined *CYCD7;1* mRNA levels in a strong *flp* allele and in the double *flp myb88* mutant (Fig. 2A). To compensate for possible differential contribution of cell types in each background, *CYCD7;1* transcript levels were normalized against genes that are (1) either ubiquitously expressed (*ACTIN2*), (2) specific for GMCs and young GCs (*FAMA*), or (3) for mature GCs (*KAT1*). *CYCD7;1* mRNA levels were elevated in both *flp* and *flp myb88* mutants (Lai et al., 2005), regardless of normalization (Fig. 2A). Consistent with the proposed negative regulation of *CYCD7;1* by FLP, we observed elevated *CYCD7;1_pro:_∷GFP* fluorescence in *flp* GMCs (Fig. 2B-C).

**Figure 2.**
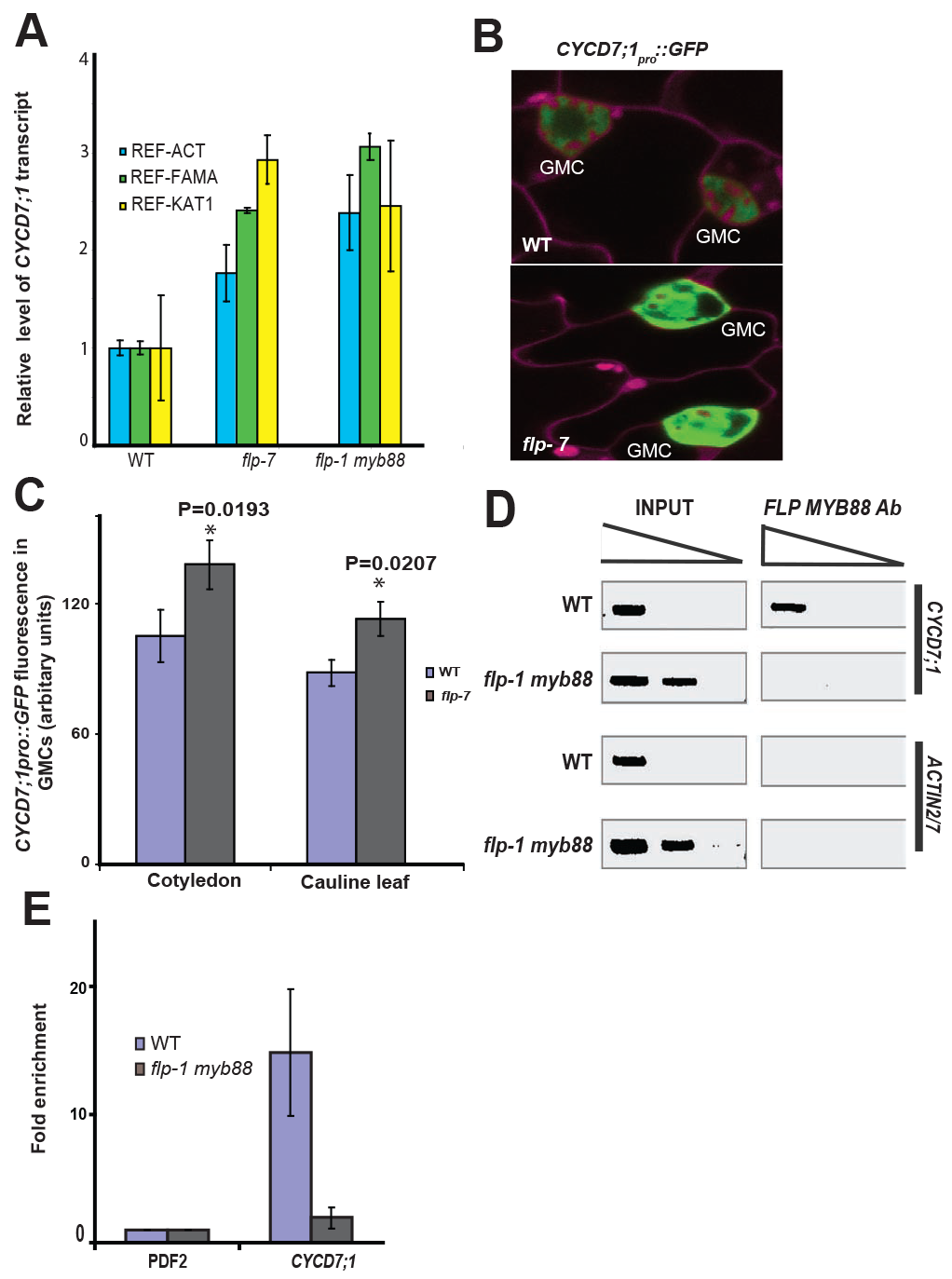
FLP moderates *CYCD7;1* transcription **(A)** Expression of *CYCD7;1* determined by qRT-PCR in WT, *flp-7* and *flp-1 myb88* normalised to *ACTIN2* (blue)*, FAMA* (green) or *KAT1* (yellow). **(B)** Expression of *CYCD7;1pro∷GFP* reporter in WT and *flp-7* mutant **(C)** Quantification of *CYCD7;1pro∷GFP* fluorescence in WT and *flp-7* GMCs in cotelydons and leaves. **(D)** PCR amplification of immunoprecipitated chromatin (ChIP) using control serum (left) and FLP/MYB antiserum (right) from WT and *flp-1 myb88* leaves. Top rows: PCR using *CYCD7;1pro* primers; bottom rows *ACTIN2/7pro* primers. Sequential dilutions from left to right. Note ChIP band only in WT extract using *CYCD7;1pro* primers (top right). **(E)** Quantification of ChIP of control (*PDF2*) and *CYCD7;1* promoter fragments in WT (blue) and *flp-1 myb88* (grey).

We next determined whether FLP/MYB88 binds to the *CYCD7;1* promoter using chromatin immunoprecipitation. An anti-FLP/MYB88 antiserum immunoprecipitated a *CYCD7;1* promoter fragment in WT, but not in *flp myb88* control extracts (Fig. 2D,E). We conclude that in the GMC, *CYCD7;1* expression is negatively regulated by FLP/MYB88 binding.

### *CYCD7;1* stimulates proliferation of the GMC

In order to understand CYCD7;1 action in the proliferating GMC, we identified a loss-of-function insertion allele *cycd7;1-1*. First we monitored the populations of stomatal precursors and stomata in the WT and *cycd7;1-1* abaxial epidermis of cotelydons over a time course of 14 days, from germination onwards (Fig. 3D), and detected an increased number of stomatal precursors, both meristemoids and GMCs, at 7 days after germination (7DAG) and 10DAG in *cycd7;1-1*, associated with a reduction of the number of stomata formed by these times (Fig. 3D). This was correlated with sustained activity of the *CYCD7;1_pro_∷GUS* and *CDKB1;1_pro_∷GUS* (Boudolf et al., 2005) reporters in the *cycd7;1-1* epidermis (Fig. 3C). These markers are normally expressed in both mutant and WT GMC (Fig. 3A,B), and so these observations are consistent with a transitory accumulation of GMCs in developing *cycd7;1-1* cotyledons. This suggested a delay in progression through the cell cycle leading to the terminal GMC division, and in order to pinpoint CYCD7;1 action we monitored nuclear size as a proxy for cell cycle progression. Whilst meristemoids have a similar average nuclear size as a population of unexpanded epidermal pavement cells, we observed a gradual increase in nuclear size as cells transit from meristemoids to guard mother cells, indicating that GMC differentiation is associated with cell cycle progression. While the meristemoids nucleus increases in size prior to the bulging associated with GMC differentiation, nuclear enlargement is slower in *cycd7;1-1* mutants suggesting a delay early on in cell cycle progression, in line with the observed larger proportion of both meristemoids and GMCs in the mutant's epidermis (Fig. 4A).

**Figure 3.**
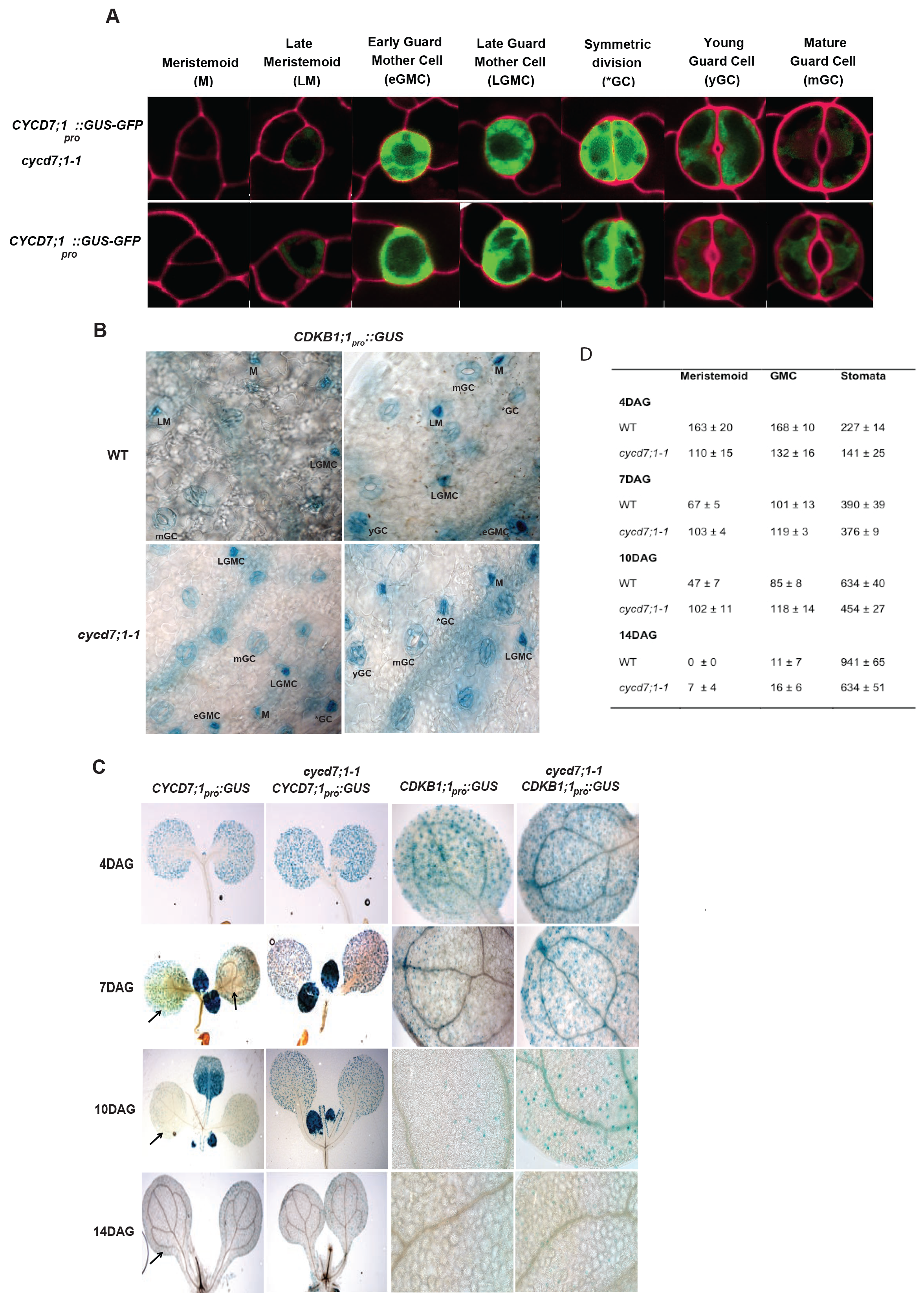
CYCD7;1 is rate limiting for the formation of stomata. **(A-B)** The expression window of the *pCYCD7;1∷GUS-GFP* and *pCDKB1;1∷GUS* markers is unaltered in the *cycd7;1-1* epidermis. **(C)** The expression of both markers persists longer in the *cycd7;1-1* epidermis. **(D)** Quantification of stomatal cell types in the abaxial epidermis of WT and *cycd7;1-1* epidermis over time reveals a prolonged presence of stomatal precursors in the mutant. The total number for each stomatal cell type in the abaxial epidermis of the cotelydon is listed.

Over time, a large proportion of *flp* mutant GCs reiterates the GMC program of symmetric division, resulting in clusters with up to 8 cells after 14 days. We observed that *CYCD7;1* expression is re-activated in these proliferating *flp* stomatal clusters (Fig. 4B) consistent with a reiteration of GMC fate. Loss of *CYCD7;1* function reduces the number of divisions of *flp* GMCs in the *flp cycd7;1-1* mutant combination (Fig. 4C). Taken together, we conclude that CYCD7;1 stimulates the cell cycle progression required for GMC division, and that its levels are moderated in the developed GMC by the FLP/MYB88 transcription factor.

**Figure 4.**
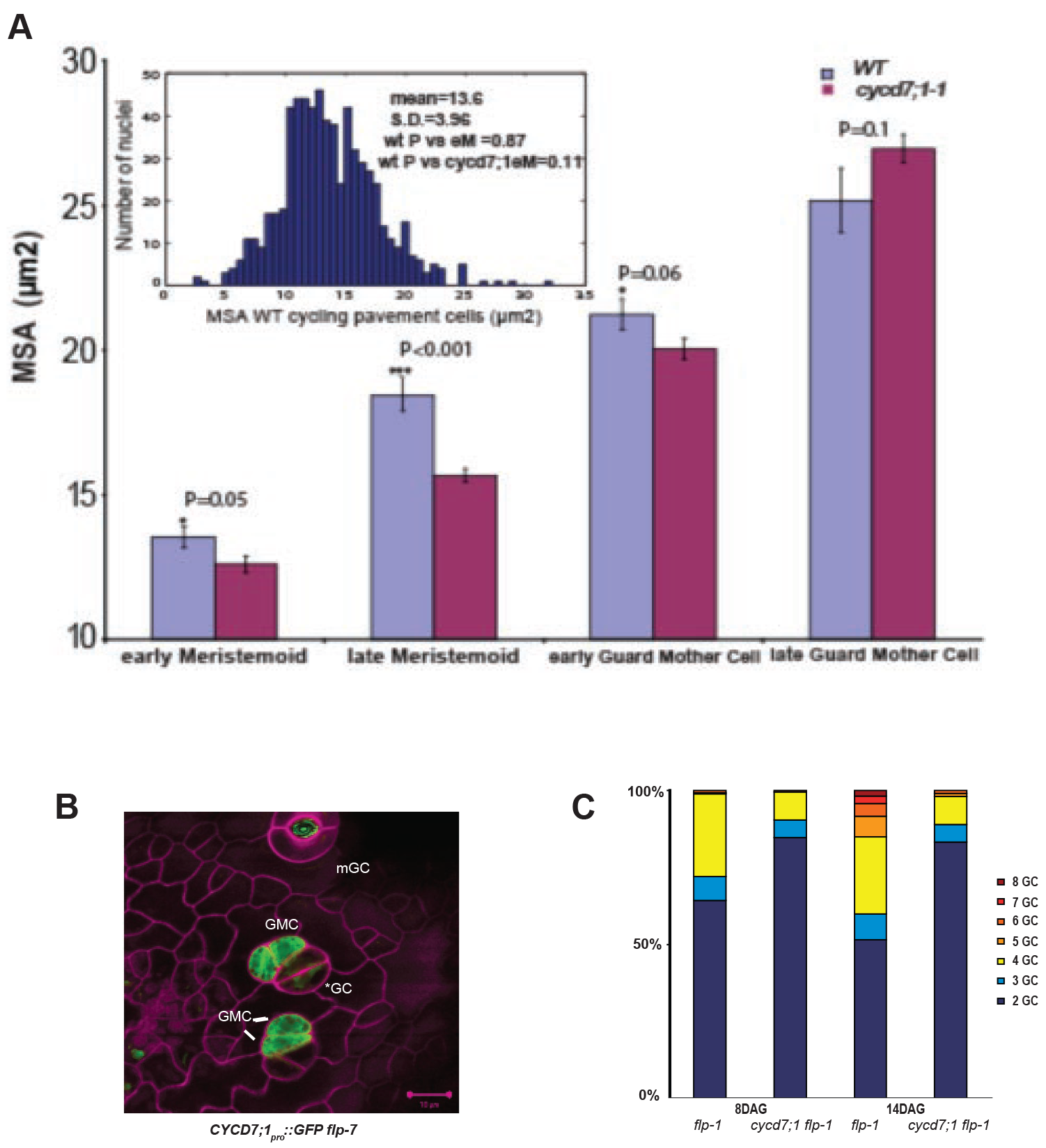
CYCD7;1 is rate limiting for the division of the stomatal precursor. **(A)** Specialisation of meristemoids into GMC is associated with nuclear enlargement. WT nuclei increase in size, reflected by the maximum surface area of a cross section (MSA) when meristemoids cells adopt the GMC morphology, a process delayed in the *cycd7;1-1* epidermis. Insert histogram depicts MSA in a population of hexagonal undifferentiated pavement cells in a young leaf (<2mm^2^). **(B,C)** Loss of *CYCD7;1* function, normally reactivated in proliferating *flp* GMCs **(B),** reduces the number of cells in *flp* stomatal clusters **(C).**

## Discussion

The formation of stomatal guard cells in the epidermis is tightly coordinated during organ growth, and requires accurate coordination of fate acquisition and cell division. Besides from generating the essential guard cells that flank and control the opening of the micropores in the epidermis, the stomatal lineage generates a large proportion of pavement cells by asymmetric divisions of meristemoids and ground lineage cells. In contrast to the asymmetric division of meristemoids, the GMC undergoes a single symmetric division upon which the newly formed guard cells become quiescent, ensuring only two cells flank the pore (Bergmann and Sack, 2007). We observed that the differentiation of a meristemoid into a bulged GMCs is accompanied with the enlargement of the nucleus, suggestive of S-phase progression. After division of the GMC, the FLP/MYB88 transcription factor limit re-entry of guard cells into a subsequent cell cycle by targeting genes encoding for core cell cycle regulators. Here we demonstrate that the *CYCD7;1* gene, encoding for a D-type cyclin expressed in the guard cell precursor, is another FLP/MYB88 target. As the loss of function phenotype for the entire *CYCD* gene family in a higher plants has not been reported, it is still unclear whether or not D-type cyclins are essential for cell cycle progression. Nevertheless, CYCDs are rate limiting for cell cycle entry in somatic tissues (Dewitte et al., 2003; Dewitte et al., 2007; Sozzani et al., 2010), as well as pathogen induced cell division (Stes et al., 2011), supporting their function in cell cycle stimulation. Our observations indicate that *CYCD7;1* stimulates the final division in the stomatal lineage and its expression is restricted by the FLP/MYB88 transcription factor.

Here we show that the transcription of different D-type cyclin encoding genes is consecutively activated in the stomatal lineage, in line with the available transcriptomics data (Adrian et al., 2015), suggesting that they are targets for pathways that mediate the different fate transitions. The Retinoblastoma-related protein (RBR), a substrate of D-type cyclin/CDK kinase complexes, interacts with FAMA, FLP and MYB88 to ensure terminal differentiation (Lee et al., 2014; Matos et al., 2014) of guard cells. RBR interacts with several transcription factors in other systems as well, and the activity of D-type cyclins is integrated in RBR mediated feedback circuits (Cruz-Ramirez et al., 2012; Sozzani et al., 2010). Now the question arises if the consecutive action of these CYCDs in the different cell types of stomatal lineage regulates the activity of the RBR interacting transcription factors by tuning the interaction with RBR.

Furthermore levels of CYCD could regulate the division potential of the different cell types. In this respect, the observation that *CYCD3;2* is a key target of the PEAPOD containing repressor complex is intriguing (Gonzalez et al., 2015).

We found that CYCD7;1 is a target of the FLP/MYB88 transcription factor, which suppress CYCD7;1 transcription, which together with the already identified targets involved in S-phase and mitosis (Xie et al., 2010) cements the function of FLP/MYB88 as an essential transcriptional repressor of core cell cycle genes for correct morphogenesis of stomata. Our observations suggest that CYCD7;1 increases the rate of cell cycle progression before the final mitosis of the GMC, in order to ensure GCs are formed timely.

## MATERIAL AND METHODS

### Plant Material

The *cycd7;1-1* (INRA-FCC172) insert line was initially generated in the Ws ecotype and has a T-DNA insertion in the second intron of *CYCD7;1,* corresponding to 715 bp downstream of the transcription start site. The allele was crosses twice into Col-0 and the absence of full-length *CYCD7;1* transcripts was confirmed by qRT-PCR using a primer set that spans the insertion site, and a primer set targeting the sequence downstream of the insertion and the levels were reduced to less than 1% of wild-type level in both cases. *CYCDpro∷GFP* and *CYCD7;1pro∷CYCD7;1-GFP* constructs were made by gateway technology and introduced into Col-O by floral dipping (Collins et al., 2012; Sanz et al., 2011). Primers are listed in supplementary table S1.

### Microscopy

Differential interference contrast light microscopy (Zeiss Imager M1) of cleared tissues was used for epidermal analysis (De Veylder et al., 2001). Confocal microscopy was performed on whole mount preparations using a Zeiss 510 or 710Meta confocal laser-scanning microscope. Cell walls were stained 1-2 mins with propidium iodide (100ug/ml, Molecular Probes).

### RNA isolation and relative transcript quantification

Relative quantification using real-time qPCR was performed using *ACTIN*, *KAT1* and *FAMA* as reference genes for normalization (Lai et al., 2005). Gene specific primers are listed (Table S1).

### ChIP of FLP/MYB88 target sequences

Antibodies against the FLP/MYB88 proteins were generated by inoculating rabbits with Ni-NTA-affinity purified NHis_6_-MYB88. ChIP experiments were performed as described (Morohashi et al., 2007; Xie and Grotewold, 2008), with modifications. Ten-day old seedlings of wild type, *flp-1 myb88* (200 mg fresh weight for each) were cross-linked in 1% formaldehyde for 20 minutes by vacuum filtration, and the cross-linking reaction was stopped by the addition of 0.1 M glycine (final concentration). Tissues were ground fine using mortar and pestle in liquid nitrogen and then suspended in 300 μl of lysis buffer (50mM HEPES, pH7.5; 150 mM NaCl; 1 mM EDTA, pH8.0; 1% Triton X-100; 0.1% sodium deoxycholate; 0.1% SDS; 1 mM phenylmethanesulphonylfluoride [PMSF]); 10 mM sodium butyrate; 1X protein protease inhibitor Sigma), and sonicated to achieve a DNA size of 0.3-1 kb. The sonication conditions using the Bioruptor (Diagenode) were as follows: At high power; 30 sec sonication followed by 30 sec break; change ice every ten minutes; 30 minutes in total. After clearing using 30 μl salmon sperm DNA/protein-A agarose (Upstate) at 4°C for at least one hour, the supernatant fractions were incubated with 1μl FLP/MYB88 rabbit polyclonal antibody at 4°C overnight. At the same time, 10% of the supernatant was saved as the input fraction. The chromatin-antibody complex was incubated with salmon sperm DNA/protein-A agarose (Upstate) at 4°C for at least 3 hours, washed with lysis buffer, LNDET buffer (0.25 M lithium chloride; 1% NP40; 1% sodium deoxycholate and 1 mM EDTA, pH8.0) and TE buffer (10 mM Tris-Cl, pH7.5; 1 mM EDTA, pH8.0) twice respectively, and the complex was reverse cross-linked in elution buffer (1% SDS; 0.1 M NaHCO_3_; 1 mg/ml proteinase K) overnight at 65°C. DNA was extracted using the PCR Cleaning Kit (Qiagen). Three biological replicates have been conducted. The ChIP-qPCR primers used for *CYCD7;1* were CYCD7;1-qP-F: 5’- AGAATGCATTTACCGCGTTT-3’ and CYCD7;1-qP- R: 5’- AAAGAAAATATGGAAGCGAGGA-3’, for *PDF2* were PDF2-F: 5’-GACGATTCTTCGTGCAGTATCGCTT-3’ and PDF2-R: 5’-GATACGGCCATGCTTGGTGGAGCTA-3’.

### Measurement of nuclear maximum cross-sectional area

Seedlings were fixed and DNA was stained in Vectashield solution with 4’,6’-diamindino-2- phenylindole (Vector Laboratories, Burlingame, CA). The GCs were examined by confocal microscopy (Zeiss meta 710). In the optical section with the largest nuclear area, the nuclear outline was traced and the MSA and average fluorescence was scored.

### Statistical tests

Due to lack of normality, non-parametric two-tailed Mann-Whitney U tests were employed.

## Acknowledgements

The authors wish to acknowledge the invaluable help from Fred Sack. We thank A. Marchbank, S. Howroyd, J. Kilby and G. Aletti for technical assistance, S. Scofield for help and discussions.

